# Lysophosphatidic acid provokes fibroblast chemotaxis through combinatorial regulation of myosin II

**DOI:** 10.1101/355610

**Authors:** Sreeja B. Asokan, Heath E. Johnson, John Sondek, Maria S. Shutova, Tatyana M. Svitkina, Jason M. Haugh, James E. Bear

## Abstract

Lysophophatidic acid (LPA), a biologically active phospholipid that is ubiquitously present in tissues and organs, provokes cellular responses such as proliferation, apoptosis, differentiation and migration via activation of G-protein coupled receptors. These receptors activate a broad range of intracellular signaling cascades to mediate these responses. Using microfluidic chambers that generate and maintain stable gradients, we observed that chemotaxis of fibroblasts to LPA has higher directional fidelity than chemotaxis provoked by the receptor tyrosine kinase (RTK) ligand platelet-derived growth factor (PDGF). Unlike fast moving amoeboid cells, mesenchymal cells such as fibroblasts do not require PI3K for chemotaxis to a GPCR ligand. In addition, the Arp2/3 complex is not required for fibroblast GPCR-based chemotaxis in either 2D or 3D environments. Our data indicate that combinatorial regulation of myosin II involving global activation by RhoA/ROCK and local inhibition of myosin II at the leading edge by PKC results in highly efficient chemotaxis of fibroblasts to LPA. Based on these observations, we develop a simple mathematical model to explain how dual regulation of myosin II is responsible for enhanced chemotaxis in LPA gradients relative to PDGF. Using pharmacological approaches, we test predictions of this model and modulate the fidelity of LPA and PDGF chemotaxis.

## INTRODUCTION

Chemotaxis or the directed migration of cells in response to soluble guidance cues is central to many physiological and pathological processes (Swaney *et al*., 2010). Cells of the innate immune system must migrate to sites of infection to effectively combat it and adaptive immune cells must migrate into, out of and within tissue compartments in response to chemokine cues to interact with appropriate target cells (Campbell *et al*., 2003). During morphogenesis, individual cells and groups of cells respond to chemotactic cues to build tissues and organs (Rogers and Schier, 2011). Upon wounding, several cell types are mobilized to the site of injury by chemotactic cues (Martin, 1997; Singer and Clark, 1999). Finally, pathophysiological processes such as cancer metastasis and fibrosis are characterized by inappropriate chemotactic cell migration (Wynn, 2008; Roussos *et al*., 2011). Thus, a more complete understanding of chemotaxis is critical for understanding and treating human diseases.

The chemotaxis of amoeboid cells is perhaps the most well studied form of eukaryotic chemotaxis and has been intensively investigated in the social amoeba *Dictyostelilum discoideum*. Under stress of starvation, *Dictyostelium* cells secrete cyclic adenosine mono phosphate (cAMP) that acts as a chemoattractant for neighboring cells (Konijn and Jastorff, 1973). The binding of cAMP to 7-transmembrane, G-protein coupled receptors (GPCRs) triggers an array of biochemical events via heterotrimeric G-proteins including activation pathways such as Ca^++^, PI3K, MAPK, Ras, and the TOR complex (Parent and Devreotes, 1999; Weiner, 2002; Swaney *et al*., 2010). Similar pathways are activated downstream of GPCRs in the leukocytes of vertebrates. Current genetic evidence suggests that some degree of redundancy exists between these pathways and/or that they act as an ensemble to transduce chemotactic signals to the motility machinery. Collectively, activation of these signaling networks lead to changes in the actin cytoskeleton, resulting in asymmetric application of force, ultimately biasing translocation of the cell towards the cue. Most studies in amoeboid cells have focused on the signaling pathways leading to generation of dendritic actin networks at the leading edge through activation of the actin branch-generating Arp2/3 complex (Rougerie *et al*., 2013).

Many non-amoeboid cell types also perform chemotaxis in various contexts. Endothelial and epithelial cells respond to chemotactic cues, often in clusters, to execute physiologic functions such as angiogenesis in the case of endothelial cells and maintaining barrier function in the case of epithelial cells. Neurons must guide subcellular processes such as growth cones to innervate targets via chemotactic mechanisms. Mesenchymal cells such as fibroblasts execute tissue repair functions such as during wound healing through chemotactic migration. In many of these non-amoeboid chemotactic scenarios, the cells are being guided by receptors of the receptor tyrosine kinase (RTK) class. Much less is known about the mechanisms of chemotaxis in these cell types, although it has long been assumed that they would use similar pathways and mechanisms as amoeboid cells (Webb *et al*., 1996; Dormann and Weijer, 2006). Therefore, it was surprising that fibroblasts depleted of two subunits of the Arp2/3 complex were capable of chemotaxis to either platelet derived growth factor-BB (PDGF) or epidermal growth factor (EGF), albeit with reduced translocation speeds (Wu *et al*., 2012; Wu *et al*., 2013). Although initially controversial, recent data suggest that neutrophil-like mammalian cells (HL-60s) and primary dendritic cells do not require the Arp2/3 complex for chemotactic migration (Collins *et al*., 2015; Vargas *et al*., 2016). Moreover, recent data from *Dictyostelium* show that the Arp2/3-activating proteins SCAR or WASP are required for efficient migration, but that they are not required for accomplishing chemotaxis (Davidson *et al*., 2018).

These observations beg the question: what force-generating cytoskeletal pathway is required for chemotaxis? Our lab recently elucidated the pathway used by fibroblasts for chemotaxis to the RTK ligand PDGF, which involves the local inhibition of non-muscle myosin II (NMII) (Asokan *et al*., 2014). During PDGF chemotaxis, the regulatory light chain of NMII is phosphorylated at Ser1/2 to inactivate or inhibit myosin function by an unknown mechanism. The kinase responsible for the phosphorylation is the classical PKC isoform PKCα, which in turn functions downstream of the second messenger diacylglycerol (DAG). Local production of DAG arises from recruitment and activation of PLC*γ*, which binds phosphorylated PDGF receptors. This pathway of NMII regulation was known in the literature (Bengur *et al*., 1987; Ikebe *et al*., 1987; Komatsu and Ikebe, 2007), but was not previously associated with direction sensing or chemotaxis. Considering that PDGF is sensed by a RTK in mesenchymal cells, whereas chemotactic signaling is mediated by GPCRs in amoeboid and other cell types, an unanswered question arising from this work was how the RTK pathway compares to chemotaxis provoked by activation of a GPCR in the same cell context.

In this work, we identify lysophosphatidic acid (LPA) as a robust, GPCR-based chemoattractant for fibroblasts. LPA, a bioactive phospholipid binds to a variety of GPCRs to trigger multiple cellular processes such as survival, differentiation, embryonic development and migration (Aikawa *et al*., 2015). A major constituent of serum, LPA has been implicated in initiation and progression of malignant diseases and is a potent chemoattractant to different cell types including tumor cells (Muinonen-Martin *et al*., 2014; Juin *et al*., 2019). Upon stimulation, LPA receptors couple to multiple heterotrimeric G proteins (Gi, Gq, G12/13 alpha subunits) to elicit cellular responses, but the specific pathway that triggers chemotaxis in fibroblasts is unclear (Moolenaar *et al*., 1997; Mills and Moolenaar, 2003).

## RESULTS

### LPA provokes a robust chemotactic response in fibroblasts via an Arp2/3-independent pathway

To investigate the mechanism of GPCR chemotaxis in fibroblasts, we observed cell migration in a gradient of LPA using a previously established microfluidic chamber system (Wu *et al*., 2012). In this chamber, mouse embryonic fibroblasts (IA32s (Cai *et al*., 2008)) that were serum starved for 3 hours prior to exposure to a gradient of LPA (4µm source concentration) were imaged overnight and tracked to measure migration speed and directionality. Forward Migration Index (FMI), net distance moved in the direction of the gradient divided by total path length, was used to quantify directionality. For this study, we considered an average FMI > 0.1 with the 95% confidence interval (CI) <u>not</u> containing zero to indicate positive chemotaxis, and experiments where the average FMI with 95% CI encompassed zero to indicate no chemotaxis. Directionality can also be represented graphically as a circular, wind-rose histogram, where the length of each leaflet corresponds to the percentage of cells migrating in that direction (where zero degrees is the direction of the gradient). The velocity of cell migration for each experimental condition was also calculated and tabulated in Table S1. IA32 cells chemotax to LPA with a high fidelity (Mov. 1) and this response is inhibited by LPAR antagonist, Ki16425 (Mov. 2) (Ohta *et al*., 2003). These data are represented graphically as a wind rose plot (Fig. 1A) and the average FMI value plotted with 95% CI (Fig. 1B). Throughout the paper, experimental conditions in FMI graphs are colored green to indicate positive chemotaxis and red to indicate no chemotaxis. Interestingly, directional migration response to 10% serum used as a chemoattractant was also inhibited by the LPAR antagonist, indicating that LPA is the main chemoattractant present in fetal calf serum (Fig. 1C).

**Figure 1:**
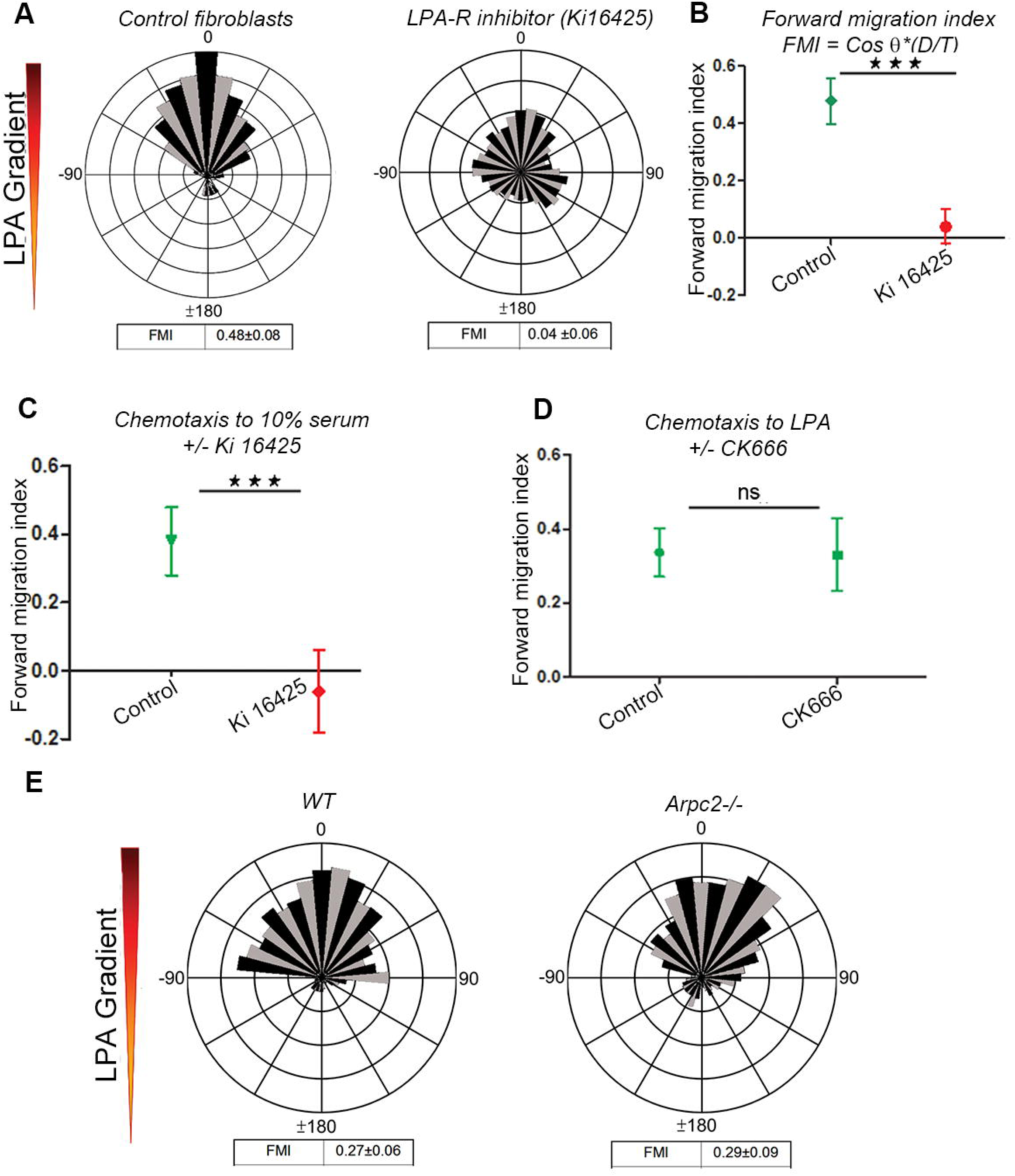
Chemotaxis to LPA, a GPCR ligand, is Arp2/3-independent. A. Cell tracks of IA32 fibroblasts chemotaxing to LPA represented as a wind-rose plot (number of cell tracked. Each leaflet represents the count frequency of cells migrating in the corresponding angular bin. Forward Migration Index (FMI) and velocity values are indicated below the plot (± 95% C.I.). Wind-rose plot showing inhibition of chemotaxis to LPA in the presence of a uniform concentration of LPARi, Ki16425 (10μM). B. Chemotactic fidelity is represented as forward migration index with 95% CI values. FMI values higher than zero with CI values that do not encompass zero are considered positive for chemotaxis. FMI range that encompass zero are considered null for directional migration. Student’s t-test was used to compare means of FMI between two data sets (*p<0.05, **p<0.01, ***p<0.001) C. 10% serum acts as a chemoattractant to fibroblasts. The LPA-R antagonist Ki16425 inhibits chemotaxis to serum. D. Fibroblasts chemotax to LPA in the presence of CK666 an Arp2/3 inhibitor with the same chemotactic fidelity as control cells E. Wind rose plots of WT (pre-Cre deletion) mouse dermal fibroblasts and *Arpc2*^-/-^ (post-Cre deletion)

Previously, we showed that the Arp2/3 complex is not required for chemotaxis to the RTK ligand PDGF. In order to examine the role of the Arp2/3 complex in GPCR-based chemotaxis, we used a chemical inhibitor of Arp2/3 (CK666). The FMI of LPA chemotaxis did not change significantly in the presence of CK666 (Fig 1D). The *Arpc2* gene encodes the essential p34 subunit of the Arp2/3 complex. Using fibroblasts derived from *Arpc2* conditional knockout mice (Rotty *et al*., 2015), we observed that genetic deletion of *Arpc2*, which leads to complete loss of the Arp2/3 complex, also does not block LPA chemotaxis (Mov. 3, 4) (Fig. 1E). Thus, LPA chemotaxis of fibroblasts, like PDGF chemotaxis, is not dependent on the Arp2/3 complex.

### Regulation of myosin IIA via myo-RLC Ser1/Ser2 is required for chemotaxis to LPA

In many systems, NMII is crucial for directional migration (Devreotes and Horwitz, 2015). To test the role of NMII in LPA chemotaxis, we treated cells with blebbistatin (BLB; a myosin II inhibitor) and observed that this treatment blocks chemotaxis in a reversible manner (Fig. 2A). Fibroblasts express the two major NMII isoforms, myosin IIA and myosin IIB, both of which are inhibited by BLB (Limouze *et al*., 2004). Although myosin IIA and IIB perform similar molecular functions, specific depletion of these isoforms result in distinct migratory phenotypes (Sandquist and Means, 2008). To assess how the different NMII isoforms are involved in chemotaxis, we used RNAi to deplete them individually. Depletion of myosin IIB did not affect chemotaxis, whereas depletion of myosin IIA ablated LPA chemotaxis in fibroblasts (Fig. 2B). Multiple labs have shown that NMIIA localizes along actin arcs and NMIIB is found on dorsal stress fibers (Burnette *et al*., 2014; Shutova *et al*., 2017; Kuragano *et al*., 2018). Immunofluorescence staining of cells during chemotaxis shows that NMIIA is distributed throughout the cell whereas NMIIB is localized towards the center of the cell (Fig. 2C). Taken together, our data show that myosin IIA is indispensable for chemotaxis and the differential localization of the two isoforms may contribute to the distinct role of myosin IIA in chemotactic migration.

**Figure 2:**
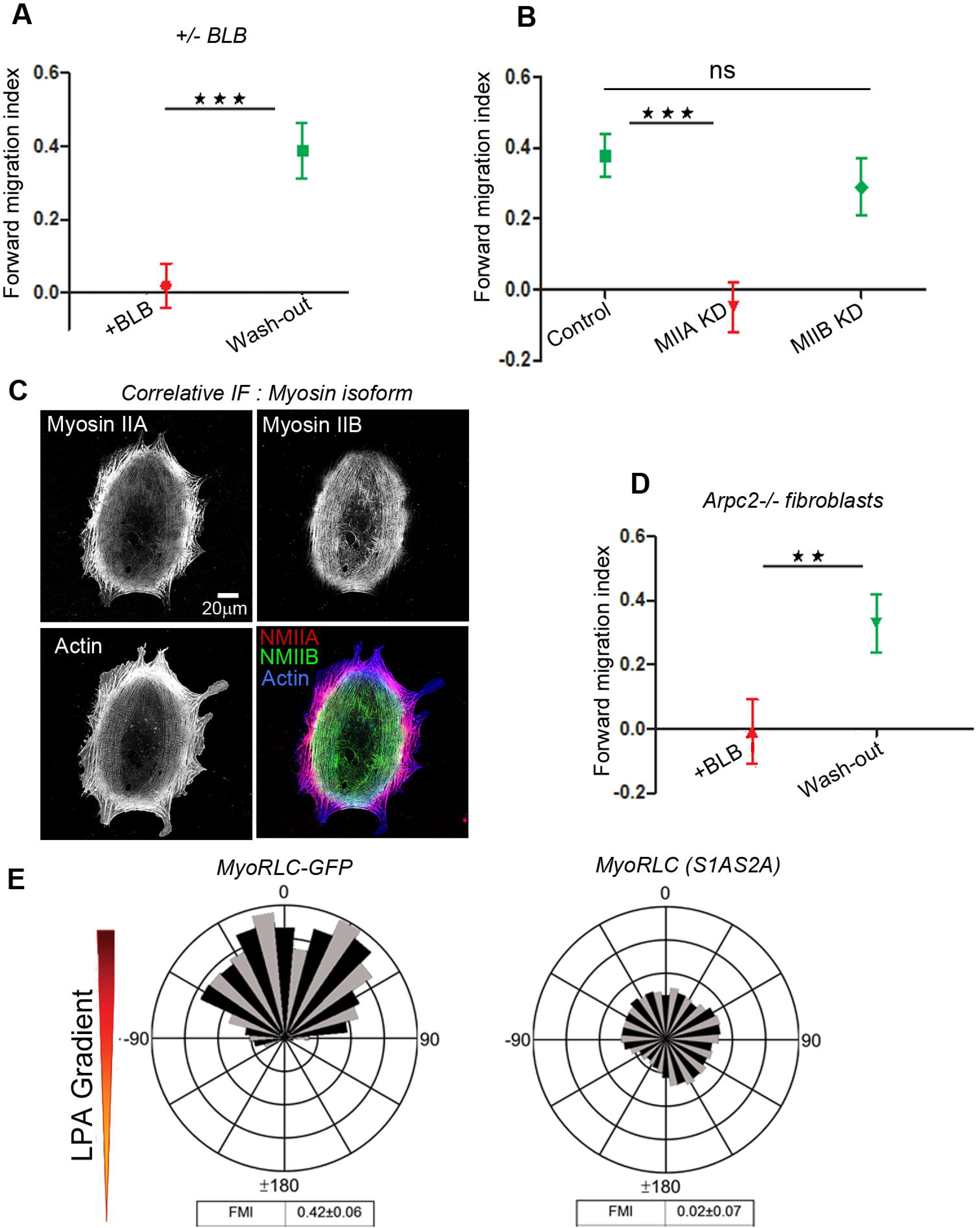
Myosin IIA isoform drives chemotaxis and is regulated by the Ser1/2 site of the myosin regulatory light chain. A. Graph of FMI shows loss of chemotaxis in the presence of blebbistatin (BLB, 15 μM) and recovery of chemotactic migration after wash-out of BLB. B. Depletion of myosin IIA using siRNA blocks cellular chemotaxis to LPA, but chemotaxis remains intact after depletion of myosin IIB C. Correlative Immunofluorescence of IA32 fibroblasts in a chemotaxis chamber shows localization of myosin IIA and Myosin IIB in chemotaxing cells. D. Calculated FMI shows *Arpc2*^-/-^ fibroblasts are unable to chemotax in the presence of blebbistatin but resume chemotaxis when the drug is washed out. E. Wind-rose plots showing chemotaxing control myo-RLC-GFP cells and non-chemotaxing mutant myo-RLC(S1AS2A)-GFP cells in an LPA gradient.

Although the Arp2/3 complex is dispensable for chemotaxis to LPA in fibroblasts, one question that arises is whether cells without Arp2/3 complex function adopt some new, non-physiological mechanism of chemotaxis. To test whether NMII is still required for chemotaxis of Arp2/3-deficient cells, we treated *Arpc2* null and control cells with BLB during LPA chemotaxis. In the presence of BLB, *Arpc2* null cells were unable to chemotax to an LPA gradient but resumed directional migration when the inhibitor was washed out and gradient restored (Fig. 2D). Therefore, NMII plays a key role in the chemotaxis of both wild-type and *Arpc2* null cells to the GPCR ligand LPA.

But how is myosin II regulated during LPA chemotaxis? The canonical pathway of activating NMII is via phosphorylation of the Thr18, Ser19 site of the regulatory light chain (RLC; also known as MLC-20, myo-RLC) by several kinases (Vicente-Manzanares *et al*., 2009). A lesser known regulatory pathway is an inhibitory phosphorylation of RLC at serines 1 and 2 (Ser1/2) (Komatsu and Ikebe, 2007; Liu *et al*., 2013). We previously demonstrated that the Ser1/2 regulatory site is required for PDGF chemotaxis by expressing GFP fusions of either the wild-type RLC or an RLC mutant containing non-phosphorylatable alanine substitutions (S1AS2A) (Asokan *et al*., 2014). To evaluate if LPA chemotaxis also requires regulation of the Ser1/2 site, we measured the chemotaxis of cells expressing wild-type (myo-RLC) or non-phosphorylatable mutants (myo-RLC S1AS2A) in LPA gradients. Consistent with previous results, expression of the S1AS2A dominant-negative RLC blocked LPA-mediated chemotaxis of fibroblasts, whereas the wild-type RLC construct did not affect this process (Fig. 2E, Mov. 5). These data indicate that regulation of NMIIA via Ser1/2 RLC inhibitory phosphorylation is critical for LPA chemotaxis.

### Chemotaxis in 3D is dependent on myosin IIA but does not require the Arp2/3 complex

Although much has been learned about the mechanism of chemotaxis by observing cells in a 2D environment, questions still remain as to how relevant these observations are to the more physiological migration of cells in a 3D environment. To address this, we polymerized a 3D collagen gel containing cells in our microfluidic chamber (Fig. 3A). Fluidic connections were made to a dual syringe pump to establish a gradient as previously described for 2D chemotaxis. Fluorescent dextran was used to monitor the gradient formed in the chamber at two different Z planes (Z = 0µm and Z = 100µm). Line scans of images of the fluorescent dextran show that the gradient is approximately the same at the two Z depths within the chamber (Fig. 3B). In the 3D environment, cells form long protruding extensions in the front and maintain a rounded morphology in the rear as seen in cells expressing myo-RLC-GFP (Fig. 3C). To investigate the role of the Arp2/3 complex in 3D chemotaxis, *Arpc2* null and wild-type (pre-Cre control) fibroblasts in 3D collagen were subjected to an LPA gradient. The *Arpc2* null cells migrated directionally with an average FMI similar to that of wild-type cells, indicating that the Arp2/3 complex is not required for fibroblast chemotaxis in 3D (Fig. 3D). To assess the role of myosin II in 3D chemotaxis, we switched to smaller REF52 fibroblasts cells that were better suited for 3D migration. The cells were depleted of the myosin IIA or IIB isoforms using RNAi (shRNA) as described previously (Shutova *et al*., 2017). NMII-depleted cells along with control cells were embedded in 3D collagen in chemotaxis chambers. While control and myosin IIB-depleted cells responded directionally to an LPA gradient, myosin IIA-depleted cells were deficient for chemotaxis (Fig. 3E). Taken together, these data indicate that the key findings on the mechanisms of chemotaxis in 2D, namely that the Arp2/3 complex is dispensable and that myosin IIA is required, hold true in 3D.

**Figure 3.**
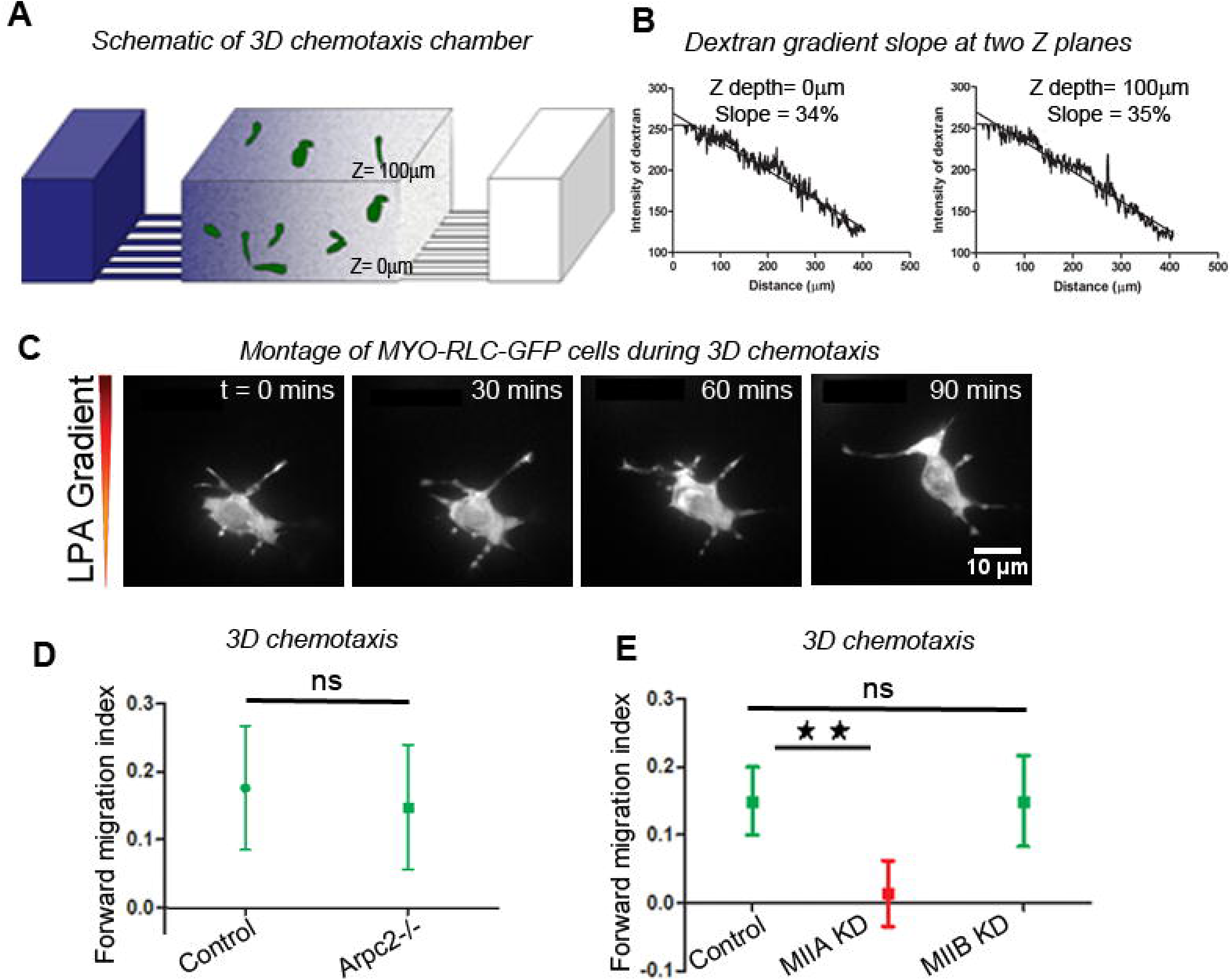
3D Chemotaxis is also driven by myosin IIA but does not require Arp2/3. A. Schematic of microfluidic chemotaxis chambers used in a 3D configuration. B. Linear intensity line scans of fluorescent dextran gradient formed in the 3D chamber calculated at Z depth of 0um and 100µm. C. Montage of REF52 cells expressing myo-RLC (GFP) chemotaxing to LPA in a 3D collagen matrix. D. FMI of WT (pre-Cre deletion) mouse dermal fibroblasts and *Arpc2*^-/-^ (post-Cre deletion) during 3D chemotaxis show Arp2/3 is not required for chemotaxis in 3D. E. FMI graph of Ref52 control fibroblasts and MIIA and MIIB KD in 3D chemotaxis chamber indicates that MIIA is required for LPA chemotaxis in 3D while MIIB is not.

### PLCβ3 and PKCα are required for LPA chemotaxis

We next set out to identify the signal transduction machinery that connects LPA receptors to NMII regulation. One logical candidate based on studies of amoeboid chemotaxis was phosphoinositide 3-kinase (PI3K) (Parent and Devreotes, 1999; Weiner *et al*., 2002); LPA receptors are thought to activate PI3K through G_i_ (Mills and Moolenaar, 2003). We performed western blotting to probe changes in p-Akt levels upon LPA stimulation and observed that the p-Akt levels did not increase with LPA stimulation but did increase upon stimulation with PDGF (Fig 4A). To functionally test the role of PI3K signaling in LPA chemotaxis, we inhibited PI3K*γ* (the relevant isoform) in LPA gradients using the inhibitor PI3-K*γ* inhibitor VII (AS041164). Treated cells were able to chemotax up LPA gradients indicating that PI3K activity is not required for mesenchymal chemotaxis (Fig 4B).

**Figure 4:**
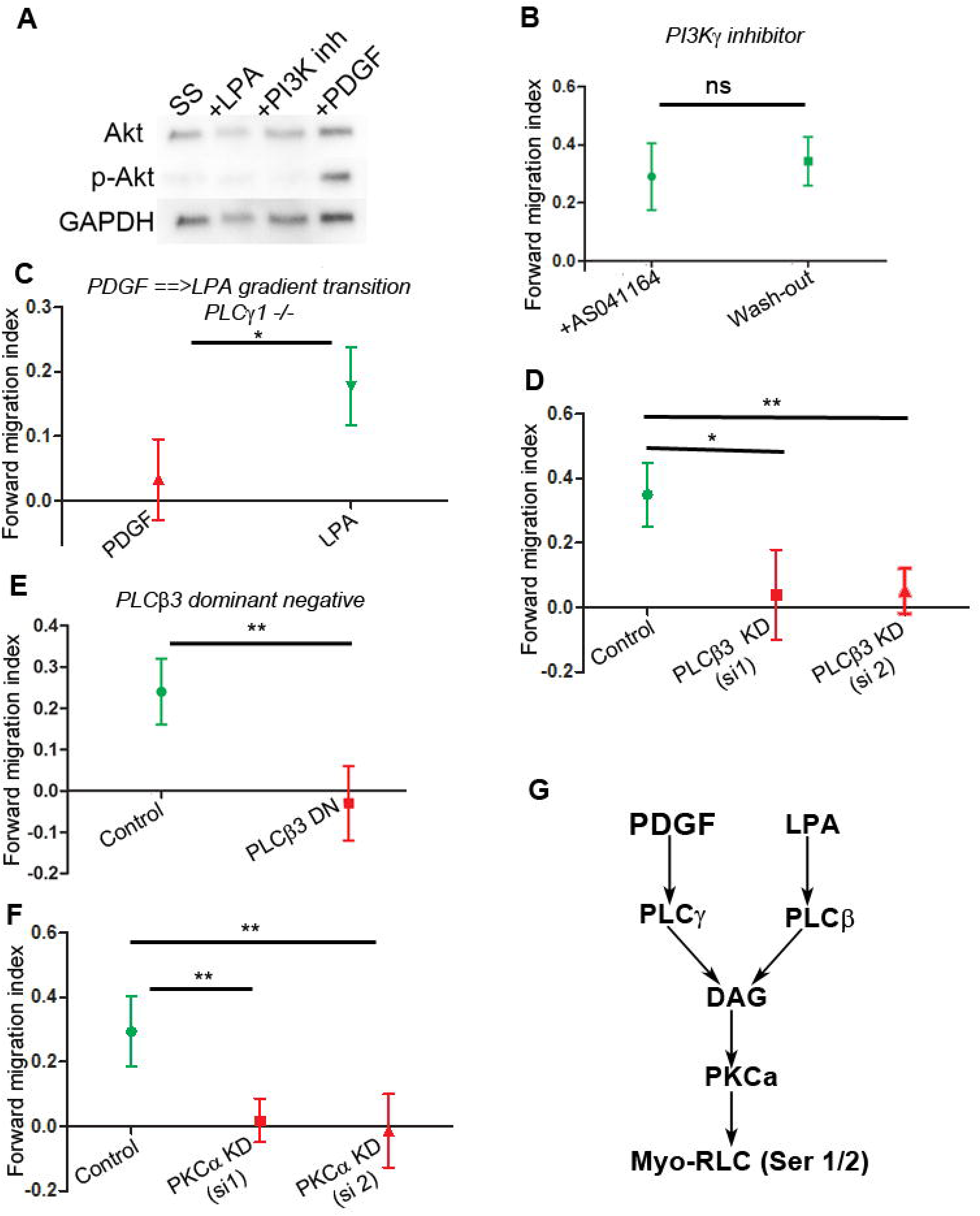
PLCβ and PKCα are necessary for LPA chemotaxis. A. Western blots probing levels of Akt and p-Akt in serum starved, LPA stimulated (2µM) and PDGF (5nM) stimulated cells. B. Cells in the presence of PI3K*γ* inhibitor can chemotax to LPA efficiently. C. PLC*γ*-null (PLC*γ*1^-/-^) cells unable to chemotax to PDGF gradients, but respond directionally to LPA gradient. D. The β3 isoform of PLC is essential for chemotaxis to LPA. E. Cells expressing a peptide inhibitor to Gαq (PLCβ3 dominant negative) are unable to chemotax to LPA. F. PKCα KD cells are unable to chemotax to LPA gradients. G. Schematic of PDGF and LPA chemotaxis pathways.

Extracellular cues like growth factors and chemokines also stimulate other pathways such as phospholipase C (PLC) (Fig 4G). We previously showed that PLC*γ*1 is required for PDGF chemotaxis (Asokan *et al*., 2014). Using PLC*γ*1-deficient fibroblasts, we initially generated PDGF gradients in our microfluidic device and observed no chemotaxis, as expected. However, halfway through the experiment, we switched the gradient source to LPA. Interestingly, PLC*γ*1-deficient cells responded directionally to the newly established LPA gradient and started to chemotax (Fig. 4C), indicating that PLC*γ*1 is not required for LPA chemotaxis. GPCRs such as the LPA receptor are known to activate PLCβ isoforms. From RNA-Seq profiling, we tested whether the most abundant PLCβ expressed in our cells, PLCβ3, was involved in LPA chemotaxis (Wu *et al*., 2013). Using multiple siRNAs, we depleted PLCβ3 and tested the cells in our LPA chemotaxis assay (Sup. Fig. 1). While control cells migrated directionally, PLCβ3-depleted cells did not chemotax to LPA (Fig. 4D), indicating that PLCβ3 is required for LPA chemotaxis. To verify this result, we also used a selective peptide inhibitor of the Gαq subunit (Charpentier *et al*., 2016). The peptide, a GFP tagged fragment of PLCβ3 harboring an I860A mutation, has a high affinity for Gαq and functions as a dominant negative for Gαq signaling, including the activation of PLCβs. Cells expressing the PLCβ3 dominant negative construct were unable to chemotax to LPA (Fig 4E).

Regardless of the isozyme, PLCs catalyze the hydrolysis of PIP_2_ into diacylglycerol (DAG) and inositol triphosphate IP_3_, the latter of which stimulates release of Ca^++^ from intracellular stores (Fig 4B) (Tanimura *et al*., 2002). Conventional PKCs are activated downstream of PLC by DAG and Ca^++^ and have a multitude of targets including serines 1 and 2 of myosin RLC. We depleted the predominant conventional PKC isoform in fibroblasts, PKCα, by RNAi and tested chemotaxis to LPA. Consistent with previous results for PDGF chemotaxis (Asokan *et al*., 2014), depletion of PKCα blocks LPA chemotaxis (Fig. 4F). Collectively, these data indicate that myosin IIA is regulated at the serine1/2 site of the myo-RLC via PLCβ3/ PKCα pathway during LPA chemotaxis (Fig. 4G).

### LPA is a more potent chemoattractant than PDGF

One observation that we made during our efforts to dissect the LPA chemotaxis pathway is that LPA is consistently a more potent chemoattractant than PDGF (Fig. 5A). Since these two factors act through similar molecular pathways, we sought to understand this difference. LPA receptors are known to couple to multiple G proteins to evoke cellular responses, and so we postulated that a signaling pathway distinct from G_q_ > PLCβ synergistically enhances the high-fidelity chemotactic response to LPA. This ligand also signals through the G_12/13_ subclass of G-proteins, which activates RhoA and its downstream kinase ROCK (Mills and Moolenaar, 2003). Stimulation of Rho-ROCK leads to the activating phosphorylation of myo-RLC (Thr18/Ser19) through a combination of direct phosphorylation by ROCK and indirectly through ROCK’s inhibition of the myosin light chain phosphatase MYPT1 (Fig. 5B) (Buhl *et al*., 1995; Fromm *et al*., 1997). Western blots of lysates from LPA-treated cells confirmed an increase in phosphorylation of MYPT1 when compared to serum starved cells (Fig. 5C). Furthermore, ratiometric analysis of immunofluorescence images revealed a significant increase in phospho-RLC/total RLC ratio in cells upon LPA stimulation (Fig. 5D). These data confirm that LPA regulates myosin II via the canonical pathway involving the activating phosphorylation of Thr18/Ser19 on myo-RLC.

**Figure 5:**
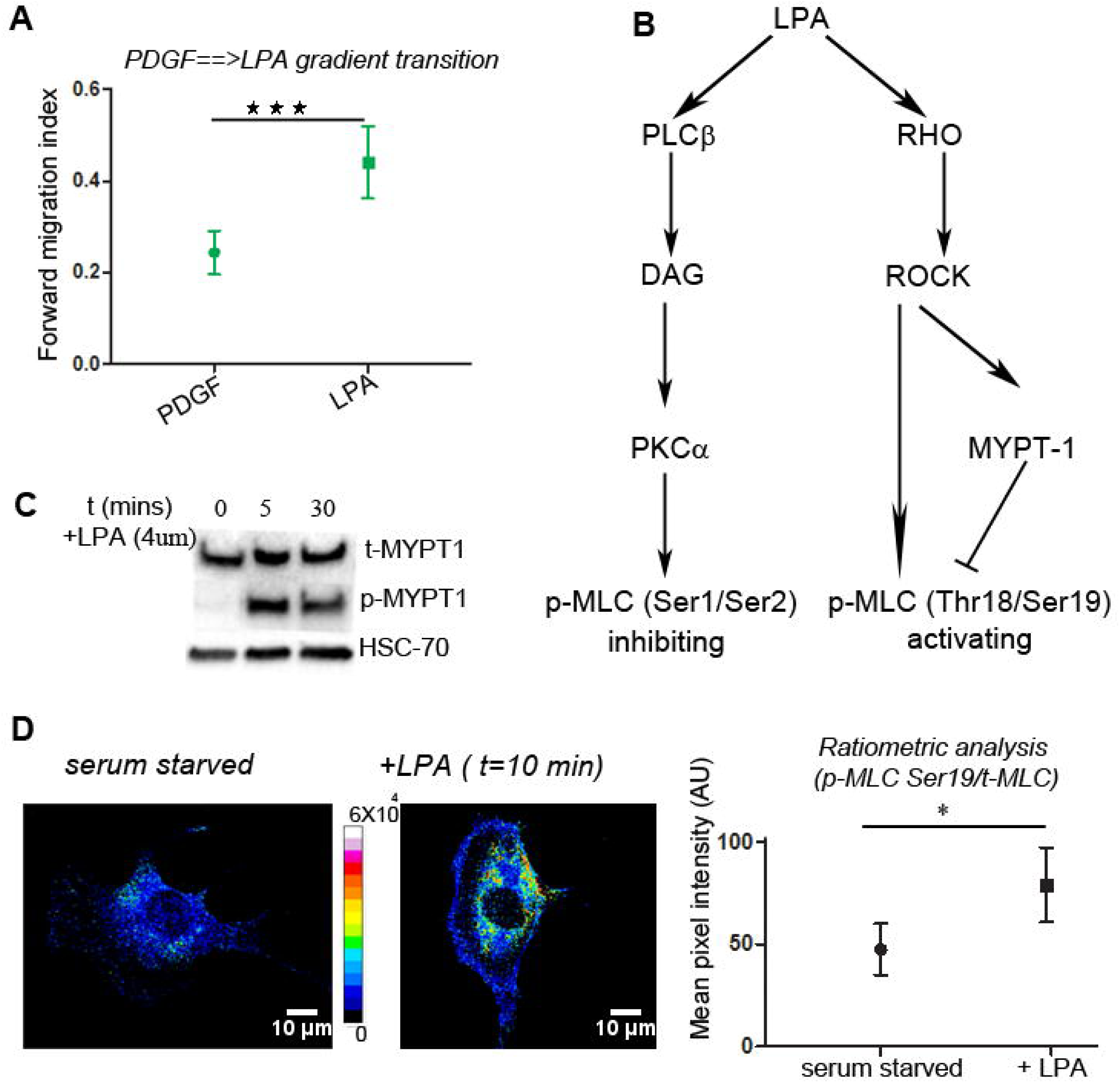
LPA is a more potent chemoattractant than PDGF. A. FMI graphs show that fidelity of LPA chemotaxis is significantly higher than that of PDGF chemotaxis. Student’s t-test was used to compare means of FMI between the 2 data sets. ***p<0.001. B. Schematic of dual regulation of myosin light chain by LPA. C. Western blots of lysates from LPA treated cells confirmed an increase in phosphorylation of MYPT1 when compared to serum starved cells. D. Histograms of pixel intensity of ratio images (p-RLC(Ser19) / t-RLC) of LPA stimulated cells show an increase in p-RLC (Ser19) when compared to serum starved cells.

### LPA gradients promote localized DAG production but global Rho activation in chemotaxing cells

Biochemical and immunofluorescence data confirmed the activation of the Rho/ROCK pathway in LPA-stimulated cells but does not provide information on spatial intracellular localization. Previously, FRET-based RhoA sensors used to study the RhoA activity profile in randomly migrating cells showed maximal activation proximal to the leading edge (Machacek *et al*., 2009). However, FRET imaging is incompatible with our current microfluidics set-up. To visualize the distribution of Rho-GTP in cells during chemotaxis, we turned to GFP-tagged RhoA binding domain of Rhotekin (RBD-GFP) (Worthylake *et al*., 2001), which has previously been used in *Xenopus* oocytes to visualize active RhoA during wound closure (Abreu-Blanco *et al*., 2014). This approach is compatible with total internal reflection fluorescence (TIRF) imaging in our microfluidic chambers. We imaged RBD-GFP translocation to the ventral plasma membrane by TIRF microscopy upon uniform stimulation with either PDGF or LPA. Analysis of fluorescence intensity shows significant increase in RBD-GFP signal after LPA stimulation, consistent with activation of RhoA (Fig. 6A). However, PDGF stimulation did not elicit a detectable increase in RBD-GFP intensity, consistent with a previous study using a FRET-based biosensor (Pertz *et al*., 2006). We then imaged cells expressing the RBD-GFP probe in an LPA gradient and observed signal throughout the cells, with only a gradual increase in signal toward the up-gradient portion of the cell (Mov.6). In comparison, when we imaged DAG using a GFP-tagged fragment of PKCα containing tandem C1 domains (Oancea and Meyer, 1998), we observed hot spots of DAG at the leading edge of the cell (Mov. 7) (Fig. 6B). To quantify this observation, we performed line scans that were 10 pixels wide and 10 microns long from the edge of a hot spot inwards towards the nucleus of the cell in cells expressing either the DAG or Rho translocation sensors. The normalized fluorescence intensity of the lines averaged across cells over the duration of chemotactic migration was plotted as a function of distance from the edge for both RBD-GFP and TandemC1 expressing cells (Fig. 6B). We used a second method for examining Rho-GTP and DAG localization across many cells relative to the gradient using previously established methods (Asokan *et al*., 2014). Briefly, we extracted 10 μm of cell periphery around the cell and projected it on a linear scale (+/-180 degrees, with 0 degrees being the direction of the gradient) over time (Fig. 6C). Consistent with our other observations, the DAG signal appears to be tightly restricted spatially at the leading edge of chemotaxing cells, while Rho-GTP appears to be activated more broadly with a bias towards the front edge of the cell (Fig. 6D). These data indicate that Rho and DAG, the two membrane-associated signaling intermediates, have different spatial distributions in chemotaxing cells, despite responding to the same external gradient of LPA ligand.

**Figure 6:**
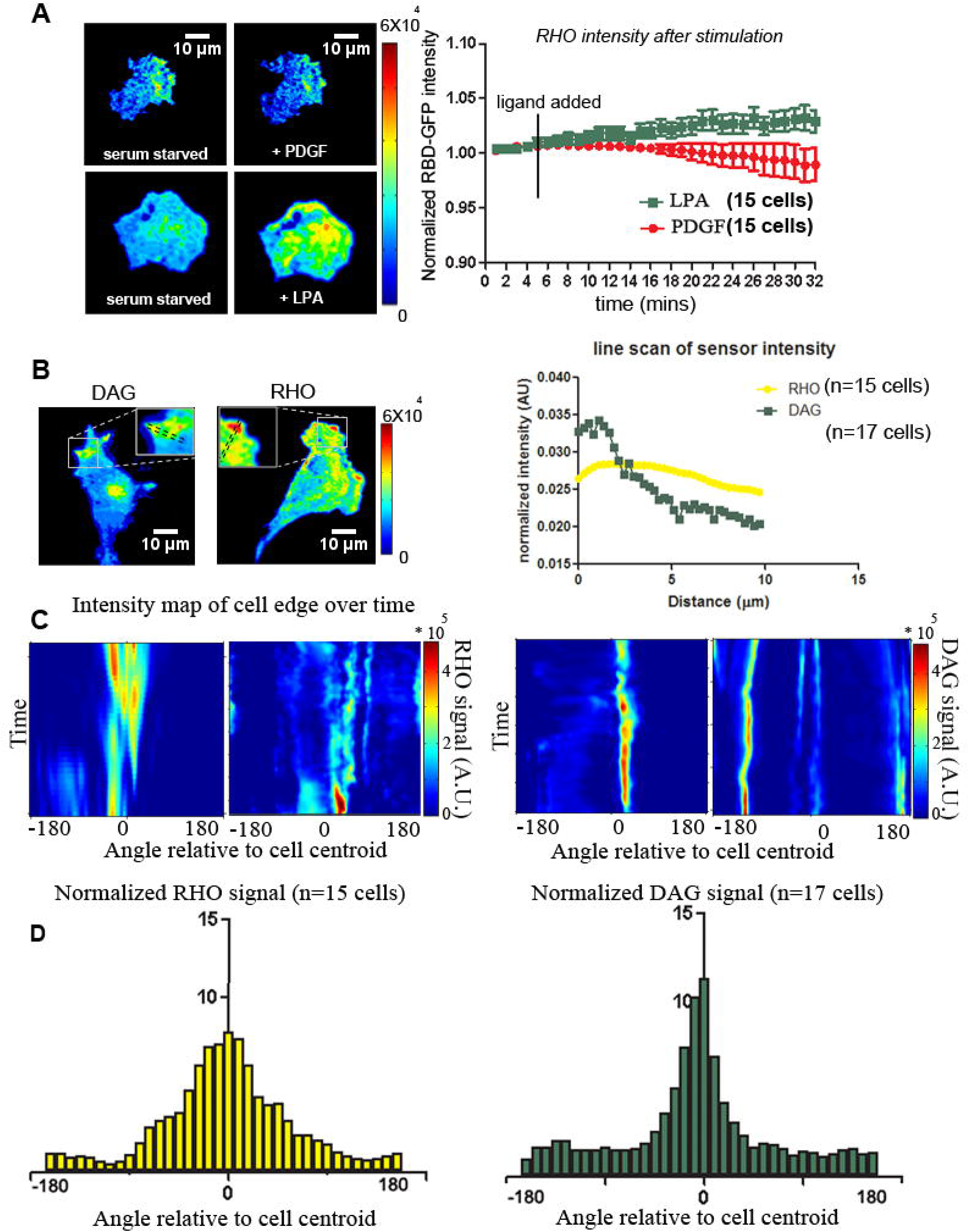
LPA invokes localized DAG activity and global Rho activity in chemotaxing cells. A. Total internal reflection fluorescence (TIRF) microscopy images of RBD-GFP localization upon uniform stimulation of cells with either PDGF or LPA. Analysis of fluorescent image intensity (mean±s.e.m.) shows significant increase in RBD-GFP signal after LPA stimulation but PDGF stimulation did not produce an increase in RBD-GFP intensity B. Line scans of RBD-GFP (Rho) and (C1)_2_-GFP (DAG) intensity at the leading edge of chemotaxing cells. C. Intensity maps of RBD-GFP and (C1)_2_-GFP expression at the periphery of cells in an LPA gradient over time. D. Histograms showing the cumulative intracellular RHO (n=15) and DAG (n=17) distribution in cells in an LPA gradient.

### Mathematical model of myosin regulation during chemotaxis predicts tuning Rho-ROCK activity affects chemotactic fidelity to LPA or PDGF

To conceptualize our experimental findings, we developed a simple mathematical model of opposing myosin regulation by PKC and ROCK. A key assumption of the model based on our experimental observations from Fig.6A is that PKC activation by DAG is far more localized than ROCK activation by Rho. Although both kinase activities are maximal at the (up-gradient) front of the cell, the relative gradient of ROCK activity is shallow, similar to that of receptor occupancy as supported by our experimental data. The other key assumption is that phosphorylation of MLC on Ser1/2 by PKC shuts off myosin activity, independent of Thr18/Ser19 phosphorylation (or, perhaps, preventing it). Thus, when both PKC and ROCK are activated during LPA chemotaxis, the localized PKC activity (solid green line) trumps the high ROCK activity (solid yellow line) at the front, yielding a substantial gradient of myosin activity that increases from front to rear (solid blue line) (Fig. 7A). In the absence of ROCK activation, as in PDGF chemotaxis, PKC alone yields a gradient of myosin activity that is predicted to be shallower (dotted blue line), which can explain the lower observed values of FMI under these conditions. The corollary of this prediction is that increased Rho-ROCK activity should increase the slope of the myosin activity gradient and consequently improve the chemotactic FMI.

**Figure 7:**
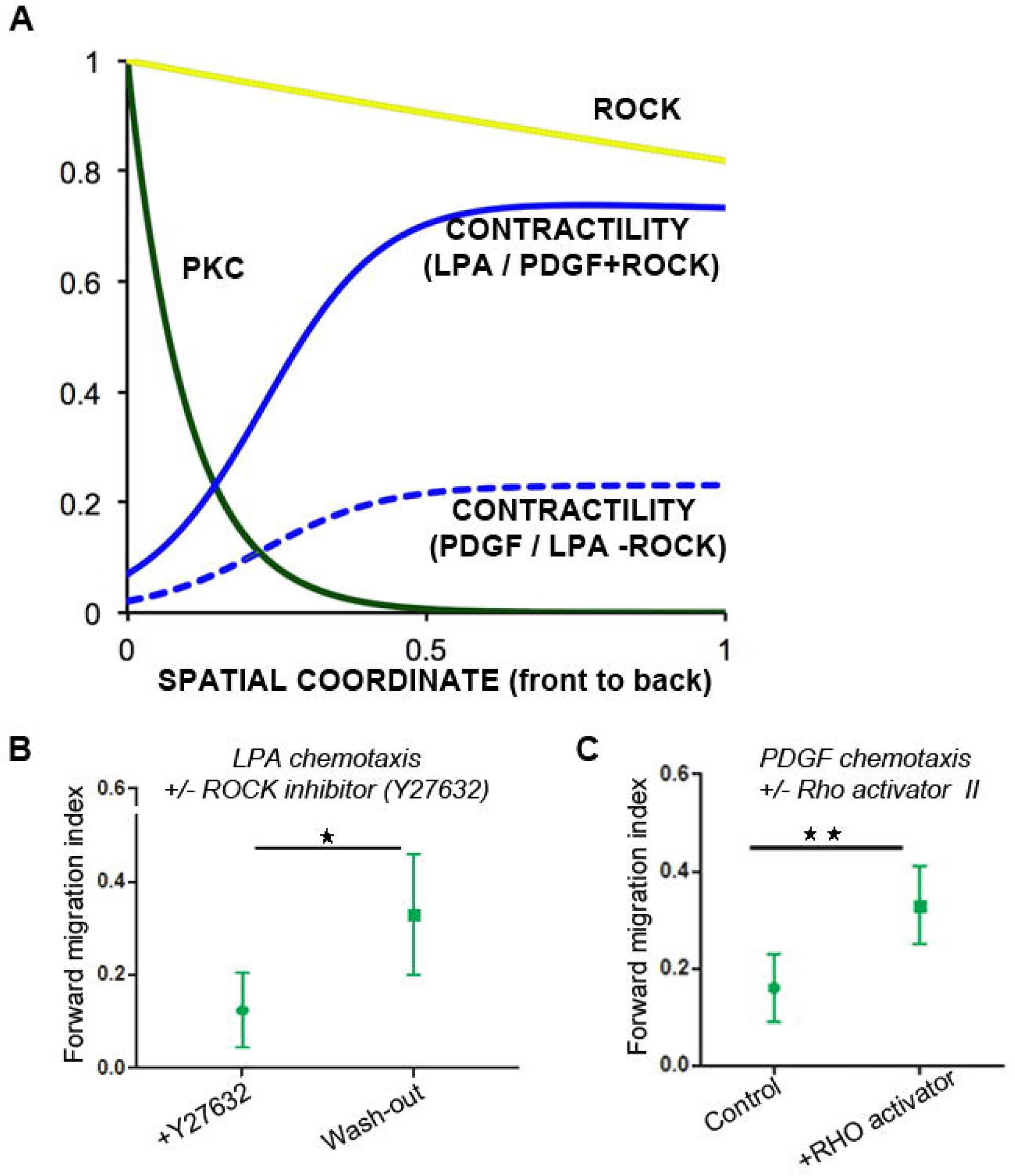
Mathematical model predicts that tuning Rho-ROCK activity affects chemotactic fidelity to LPA or PDGF. A. Graph showing expected PKC and ROCK activity from experimental observations and predicted myosin activity/contractility from our mathematical model from front to back of a chemotaxing cell. Higher asymmetric myosin activity translates to higher force generation and better chemotactic fidelity. B. Inhibition of ROCK activity by inhibitor Y-27632 during LPA chemotaxis reduces the FMI of migrating cells reversibly. When the inhibitor was washed out, the FMI increased to control values for LPA chemotaxis. In alignment with our experimental observations, our model predicts a shallow myosin contractility profile for LPA chemotaxis in the presence of ROCK inhibition (dotted blue line) and a steeper myosin contractility profile during control LPA chemotaxis (solid blue line). C. The FMI of PDGF chemotaxis increases when ROCK is activated externally by Rho activator II. In alignment with our experimental observations, our model predicts a shallow myosin contractility profile for PDGF chemotaxis (dotted blue line) and an increase in myosin contractility during Rho activation (solid blue line).

We sought to experimentally test these predictions. To assess the mechanistic role of Rho activation, we treated fibroblasts with ROCK inhibitor (Y27632) during LPA chemotaxis. This treatment produced a significant reduction in FMI that could be reversed by washing out the drug (Fig. 7B, Mov. 8), consistent with the predicted role of Rho/ROCK pathway in increasing the fidelity LPA chemotaxis. We also tested whether activating the Rho-ROCK pathway could improve the average FMI of PDGF chemotaxis. We used the Rho Activator II, a cell-permeant catalytic fragment of the bacterial CNF toxin that deamidates glutamine-63, thereby increasing the level of GTP-bound endogenous RhoA (Schmidt *et al*., 1997). Interestingly, the average FMI of PDGF chemotaxis increased significantly in the presence of Rho activator II (Fig. 7C, Mov. 9). Thus, consistent with the predictions of our model, our experimental data indicate that the Rho/ROCK pathway acts synergistically with the PLC/PKC pathway to increase the fidelity of fibroblast chemotaxis to LPA.

## DISCUSSION

In this work, we dissected the mechanism of fibroblast chemotaxis to the GPCR agonist LPA in direct observation microfluidic chambers. Similar to our earlier findings with PDGF, we established that the PLC/PKC/NMII pathway is essential for LPA chemotaxis. However, we also found some notable differences. PDGF chemotaxis utilizes the PLC*γ*1 isozyme, whereas LPA chemotaxis requires PLCβ3 to produce a spatially constrained area of DAG at the leading edge of chemotaxing cells. For both chemotactic ligands, localized DAG activates PKCα, which locally *inactivates* NMIIA via phosphorylation of the Ser1/2 site in the myo-RLC. Interestingly and distinct from PDGF, LPA also simultaneously *activates* NMII via the Rho/ROCK pathway and phosphorylation of myo-RLC on Thr18/Ser19. These two pathways functionally cooperate to regulate NMII activity and produce high fidelity chemotaxis to LPA. We developed a mathematical model to explain how the internal gradients of PKC and ROCK activities synergistically lead to a steep intracellular gradient of NMII activity that we experimentally tested by tuning the Rho-ROCK pathway during LPA and PDGF chemotaxis.

### PDGF vs. LPA chemotaxis in fibroblasts

One of our main goals was to compare and contrast chemotactic responses elicited by distinct classes of receptors—RTKs vs. GPCRs. Although both receptor classes have been extensively studied for their ability to mediate chemotactic signaling in different cell types, we know very little about how the downstream pathways differ. Using fibroblasts with endogenous expression of both receptor types, we have been able to make this comparison. Downstream of both receptors, we find that a similar molecular pathway is required, involving the inactivation of NMII by PKC phosphorylation of the regulatory light chain of NMII at Ser1/2. A critical signaling intermediate required for PKC activation is the local generation of DAG by PLC. In the case of PDGF (RTK), this is accomplished by PLC*γ*1, while in the case of LPA (GPCR) this is carried out by PLCβ3. It will be interesting in future work to consider the regulation and possible signaling feedbacks affecting the recruitment and/or enzyme kinetics of these two PLC isozymes during chemotaxis.

Two things that do not appear to be required for fibroblast chemotaxis to either PDGF (Asokan *et al*., 2014) or LPA chemotaxis (this study) are PI3K activity or the Arp2/3 complex. In the case of PI3K, early results suggested that this activity was critical for chemotaxis in *Dictyostelium* and neutrophil-like cells, but more recent findings indicate that PI3K signaling is not strictly required for chemotaxis; rather, it may only be essential when cells encounter shallow gradients of chemoattractant (Takeda *et al*., 2007; Bosgraaf *et al*., 2008). In this work, we observed that LPA does not increase p-AKT levels (a downstream PI3K target) and that fibroblasts chemotax to LPA in the presence of a PI3K*γ* inhibitor, both consistent with PI3K not being required for fibroblast chemotaxis. The dispensable nature of the Arp2/3 complex for chemotaxis has also been somewhat controversial (Suraneni *et al*., 2012), but other recent studies support this conclusion (Collins *et al*., 2015; Vargas *et al*., 2016). Since significant effort has gone into connecting chemotactic receptors to activation of the Arp2/3 complex, this result is challenging, but it demands that we consider new mechanisms of chemotaxis.

LPA is not only a useful compound to probe the mechanisms of GPCR chemotaxis in our *in vitro* system but is also an important signaling molecule in various physiological and disease-related contexts. Six LPA receptor (LPAR) genes have been identified and characterized to date: *LPAR1*-*LPAR6* in humans and *Lpar1*-*Lpar6* in mice (Kihara *et al*., 2014; Yung *et al*., 2014). These seven-transmembrane GPCRs bind different forms of LPA with varying affinities and activate specific heterotrimeric G protein pathways defined, in part, by Gα_12/13_, Gα_q/11_, Gα_i/o_ and Gα_s_. The downstream signaling cascades involve well-known mediators such as Ras, Rho, Rac, Akt, MAPK, PKC and adenylate cyclase. The resultant activation of these signaling cascades then influences major cellular processes such as proliferation, apoptosis, morphological changes, migration and differentiation. We have shown that LPA is a potent chemoattractant for fibroblasts, acting via receptors 1 and 3 that are expressed in fibroblasts. This directional response was inhibited by Ki16425, an LPA1-3 specific antagonist. Interestingly, Ki16425 also inhibited chemotaxis to serum, identifying LPA as the major chemoattractant in serum.

### Signaling pathways, downstream from the same receptor, have distinct spatiotemporal characteristics

One of the difficulties in studying chemotaxis is grappling with how complex and parallel signaling pathways cooperate to produce directed migration. Although our studies have identified an obligatory pathway for chemotaxis, our data also indicate that other signaling events contribute to the fidelity of this process. LPA, through its receptor, stimulates multiple heterotrimeric G proteins: G_q_ to activate PLC, G_i_ to activate PI3K and others and G_12/13_ to activate RhoA. Our data point to PLCβ3 as the key downstream enzyme that must be activated by G_q_ for LPA-directed migration. However, LPA activation of the Rho/ROCK pathway by G_12/13_ acts synergistically to improve chemotactic fidelity. The linear external LPA gradient is converted to internal gradients of the signaling intermediates DAG and active RhoA that display different spatial profiles. While DAG is localized and highly polarized to the front of the cell, Rho activation shows a graded bias from the front to the back of the cell that drops in a roughly linear fashion. This difference likely arises from reaction-diffusion properties of the enzymes and of their products and corresponding feedback regulation. Regulators of DAG include the DAG kinases (Merida *et al*., 2008), which convert DAG to phosphatidic acid (PA) and thus limit the spatial range of DAG diffusion in the plasma membrane. Another likely regulator of this pathway is MARCKS, a PKC substrate that sequesters PIP_2_ via its effector domain (Hartwig *et al*., 1992). We have begun to study this signaling circuit theoretically, formulating reaction-diffusion models that explore the amplification of internal DAG gradient during chemotaxis (Mohan *et al*., 2017). On the other side of the signaling network, the distribution of active RhoA is governed by guanine nucleotide exchange factors (GEFs), GTPase activating proteins (GAPs) and guanine nucleotide dissociation inhibitors (GDIs) (Marjoram *et al*., 2014). The complexity and inter-relatedness of these signaling pathways will likely require more sophisticated experimental and modeling efforts to unravel. But, how do the two signaling pathways work in parallel to produce robust chemotactic migration? We have identified myosin II as the likely downstream effector, a common target for both the DAG-PKC and RhoA-ROCK pathways.

### Generating asymmetric force distribution by regulating myosin during chemotaxis

From first principles, a key step in chemotaxis is converting the spatial bias encoded by the external gradient of chemoattractant into asymmetric force generation required for migration in the direction of gradient. Through our observations of LPA chemotaxis, we have shown that the key force generator directing migration is myosin IIA. Asymmetric myosin activity likely regulates multiple processes relevant to migration including primary effects on contractility of bundled actin structures, as well as secondary effects on processes such as focal adhesion dynamics. The canonical regulation of myosin occurs via phosphorylation of Thr18/Ser19 on RLC, which activates myosin. However, the non-canonical inactivating phosphorylation of myosin RLC at the Ser1/Ser2 site is critical for chemotaxis. During PDGF chemotaxis, DAG and membrane-bound PKC are concentrated in hot spots at the up-gradient side of the cell, selectively inactivating myosin in those regions. This occurs in a background of some basal level of myosin activity. In the case of LPA chemotaxis, RhoA is activated throughout the cell with a linear gradient from front to back, accompanied by ROCK and myosin activation. Concurrent with RhoA activation, LPA activates PLCβ3 to generate a highly concentrated area of DAG on the up-gradient side of the cell. This leads to local PKC activity and inactivation of myosin via ser1/2 phosphorylation. Based on the literature (Ikebe *et al*., 1987) and our previous data, Ser1/2 inactivating phosphorylation is dominant over Thr18/Ser19 activating phosphorylation, thus generating a relatively steep gradient of myosin activity with more active myosin towards the rear of the cell. In this model, Rho-ROCK activation serves to amplify the myosin gradient which we postulate increases chemotactic fidelity. Consistent with this idea, when the Rho activator is included during PDGF chemotaxis, we observe higher chemotactic fidelity. Conversely, inhibition of ROCK during LPA chemotaxis reduces the chemotactic fidelity. This *<u>g</u>lobal <u>e</u>xcitation* of myosin activity by ROCK and *<u>l</u>ocal <u>i</u>nactivation* by PKC (GELI) model for enhanced chemotactic migration will require further refinement but provides a conceptual framework to test new ideas about chemotaxis.

### Diversity in the mechanisms of directed migration

Perhaps the most important observation arising from this study is that cells have a variety of ways to achieve directed migration. Signaling pathways like PI3K/PTEN, TOR, Ras and PLA are collectively important for *Dictyostelium* chemotaxis, but seem less important for mesenchymal cell chemotaxis. Instead, these cells use an obligatory PLC-PKC-Myosin II pathway to achieve chemotaxis downstream of both RTKs and GPCRs. Mesenchymal cells like fibroblasts have actin stress fibers and robust focal adhesions to bind to and contract the extracellular matrix as part of their biological function in tissue repair. It is therefore not surprising that myosin II plays a central role in regulating their chemotaxis. Interestingly, these same cells use a different pathway to respond to gradients of substrate bound extracellular matrix (haptotaxis). Here, they rely on the Arp2/3 complex to produce lamellipodial protrusions that explore their local environment (King *et al*., 2016). Biased lamellipodia during haptotactic migration leads to directed whole cell migration up gradients of ECM proteins. As we begin to delineate the mechanisms of directed migration for different cell types and towards different migration cues (e.g., durotaxis, galvanotaxis), we anticipate that a diverse set of pathways and mechanisms will be revealed.

## Supporting information

Mov1

Mov2

Mov3

Mov4

Mov5

Mov6

mov7

Mov8

Mov9

## ACKNOWLEDGEMENTS

We gratefully acknowledge helpful discussions with Robert Insall, Gregg Gundersen and David Graham during the early stages of this project. This work was supported by NSF grant to S.B.A. (1706019), NIH grants to J.E.B. (GM110155), J.E.B and J.M.H (GM111557, EB018816), J.S. (GM120291, GM057391) and T.M.S. (GM095977).

## AUTHOR CONTRIBUTIONS

Conceptualization, S.B.A, J.M.H, J.E.B; Investigation, S.B.A; Formal Analysis, J.M.H; Resources, J.D.S, M.S.S, T.M.S.; Writing, S.B.A, J.M.H., J.E.B; Funding Acquisition, S.B.A, J.M.H, J.E.B.

## MATERIALS AND METHODS

### Reagents and Materials

AlexaFluor-conjugated phalloidins were from Invitrogen. Antibodies were purchased from Cell Signaling Technology (pMLC, NMIIA, NMIIB), ECM Biosciences (t-MLC) and Santa Cruz Biotechnology. LPA was purchased from Sigma. Inhibitor of LPA-R (Ki16425) was purchased from Selleckchem, PI3Kγ (Inhibitor VII) from EMD millipore, Blebbistatin and ROCK inhibitor (Y27632) from Sigma Aldrich. siRNAs were purchased as a set of four from Qiagen for myosin IIA (GS17886), myosin IIB (GS77579), PKCα (GS18750) and PLCβ3 (GS18797).

### Immunofluorescence and western blots

For correlative immunofluorescent staining, the cells were fixed and stained in microfluidic chambers as described previously (Samantha J. King *et al*., 2016). After 12-hour chemotaxis experiment, cells were fixed with 4% PFA and permeabilized in 0.1% Triton X-100 in PBS for 5 mins. Cells were then blocked for 15 mins in PBS containing 5% normal goat serum (Jackson Laboratories) and 5% fatty-acid-free BSA. Primary antibodies were applied to cells in PBS containing 1% BSA for 1 hr at room temperature. Cells were stained in various combinations with AlexaFluor-647 phalloidin for F-actin (1:200 dilution), myosin IIA (1:250 dilution) or myosin IIB (1:250 dilution) antibodies. After washing the cells three times in PBS, fluorescent dye-conjugated secondary antibodies were diluted to 1:250 in 1% BSA in PBS and flowed into chamber for 1 hr. Images were captured using a FluoView scanning confocal inverted microscope (FV1000, Olympus) equipped with a 40x objective, a Hamamatsu PMT and controlled by Fluoview software. Maximum intensity projection was determined with ImageJ from a z-stack. Western blotting of whole cell lysates was done by standard techniques. Cells were serum starved overnight and pre-treated with 1 μM LPA for the times indicated unless stated otherwise. Blots were probed using primary antibodies followed by incubation with HRP-conjugated secondary antibodies. The membrane was scanned in a ChemiDoc MP imager (Bio-Rad).

### Cell culture, viral transduction and plasmid/siRNA transfections

Cells were cultured in DMEM supplemented with 10% FBS (HyClone) and 292 μg/mL L-glutamine. Transient transfections were performed using FuGene 6 (Roche) for HEK293 FT cells and Lipofectamine 2000 (Thermo Fisher) for fibroblast cell lines. Lentivirus production, infections, and fluorescence-activated cell sorting were performed as described (Cai *et al*., 2008). PLC*γ*1^-/^, PLC*γ*1 rescue cells and peptide inhibitor to the Gαq subunit were kind gifts from Sondek lab (UNC-CH). Ref-52 cell lines with myosin IIA and myosin II B shRNA knock downs were generated and described previously (Shutova *et al*., 2017). Cells expressing tagged tandem C1 and RBD-GFP domains were generated by transient transfection protocols. Tagged tandem C1 domains of PKCδ, (C1)_2_-GFP was a gift from Tobias Meyer (Addgene plasmid #21212) and RBD-GFP from Burridge lab. Plasmids were transiently transfected into IA32 fibroblasts using Lipofectamine 2000 reagent. The cells were sorted using a Biorad S3 sorter 18 h post-transfection. The sorted cells were immediately loaded into the chemotaxis chamber for cell migration assays. myosin II, PKC and PLC knock down cells were generated using standard siRNA knockdown protocols. siRNA for each gene was purchased as a FlexiTube Gene Solution set from Qiagen and reconstituted following manufacturer’s instructions in nuclease-free water. IA32 fibroblasts were transfected at 50% confluency using 25-50 pmol of siRNA and RNAiMAX transfection reagent (Life Technologies) following the manufacturer’s protocol. Knockdown was analyzed by blotting and/or immunofluorescence 48-96 hours post-transfection. Dermal fibroblasts (26DF) were cultured in DMEM supplemented with 10% FBS and 1% Glutamax. 26DF cells were plated at low density and treated several hours later with 2 μM 4-hydroxy-tamoxifen (4-OHT) to induce recombination of the *Arpc2* allele. Two days later, fresh media was added with a second dose of 2 μM 4-OHT. On day 5, cells were harvested, lysed or expanded for subsequent experiments.

### Generation of myoRLC-GFP cell lines

pLL6-MYH12A construct was made by replacing 5’LTR promoter in pLL5.0 (Cai *et al*., 2008) with P_Tight_ promoter from pTRE-Tight Vector (Clontech) and inserting human MYH12A using EcoRI-BamHI cloning sites. MYH12A insert was obtained by PCR from cDNA isolated from human cell line WM-266-4 (ATCC CRL-1676™). Point mutations in MYH12A were introduced using standard overlap extension technique using KOD polymerase (EMD Millipore). Tetracycline-inducible MYH12A WT and S1AS2A-mutant IA32 cell lines were made by first preparing a stable rtTA-expressing IA32 cells using pLenti-CMV-rtTA3-hygro lentivirus infection (virus made with Addgene plasmid 26730) and selection with 100 μg/mL hygromycin. These stable rtTA cells were then infected with pLL6-MYH12A-GFP lentivirus, and subjected to 3 rounds of FACS: first, cells were induced with 1ug/mL Doxycycline (Dox) and sorted for GFP-positive cells. The resulting cell population was then allowed to recover to uninduced state in the absence of Dox and sorted for no GFP expression, and, finally, the cell population obtained after this sort was induced with 1 μg/mL Dox again and sorted for high GFP expression. Resulting Tet-inducible MLC populations were induced with 1 μg/mL Dox for 24 hours before chemotaxis experiments.

### Directional migration image acquisition and analysis

PDMS microfluidic devices were prepared as described previously (Wu *et al*., 2012). Cells were plated in the middle chamber of the microfluidic device in serum free media and starved for up to 3hrs prior to chemotaxis experiments. LPA (Sigma) was dissolved in a 1:1 ratio of distilled water:ethanol to generate a 1 mM stock solution and stored at −20°C. For use as a source in chemotaxis chambers, 4 μl LPA stock was added to 1ml DMEM containing 2% serum and 0.05% (w/v) BSA. DMEM containing 2% serum and 0.05% (w/v) BSA was used for the sink channel. For drug treatments, the cells were pretreated for the recommended time and the drug was also added to the source and sink tubing to maintain a steady concentration of drug. Chemotaxis assays were performed on an Olympus microscope with a 20X objective using Metamorph imaging software. Images were collected every 10 minutes for up to 24 hours. Individual cells were manually tracked using ImageJ software (Manual Tracking plugin). Only viable and visibly migrating cells (net path length > 50 μm) were tracked during the experiments. We performed a minimum of 3 independent repeats for each chemotaxis experiment to track a minimum of 30 cells per experiment. The tracks obtained were analyzed using the Chemotaxis Tool ImageJ plugin. This analysis tool was used to extract the forward migration index (FMI) along with the velocity of migration. Further, the plugin also generates histograms (count frequency) of migration direction of cells for each data set. The secplot function of Matlab was then used to generate rose plots of directional migration on normalized polar coordinates, where the outer most ring corresponds to frequency (r) of 8%. Ratio imaging:

### TIRF microscopy

The microfluidic chamber system is fully integrated with live-cell TIRF microscopy imaging. A 60X TIRF objective was used to image the translocation of (C1)_2_-GFP or localization of the RBD-GFP constructs during chemotaxis. The cells were imaged at 2-minute intervals for 5-10 hours.

### Ratiometric image analysis

Unstimulated control cells and LPA stimulated cells were fixed and stained with total RLC and phospho RLC antibodies, using routine immunofluorescence protocol. Coverslips were imaged on an FV1000 confocal microscope with a 40X, 1.3NA objective and images captured with a Hamamatsu photonics PMT. The images were analyzed using the image calculator tool in Image J. The target of interest (phosphor-MLC) was set as the numerator and the control (total-MLC) was set as the denominator. The resulting ratio image was multiplied by a mask of the thresholded denominator to reduce background noise.

### Analysis of the DAG and Rho pattern

To correlate the intensity of DAG and Rho in a cell and direction of chemotaxis, signaling maps of tandem C1 and RBD-GFP probes were created. These fluorescence intensity maps were created for each cell by a process of thresholding, masking and segmenting within angular bins relative to cell centroid (Welf *et al*., 2012). The normalized patterns are then averaged over all the cells for each condition to produce an aggregate plot. The linear (unbinned) plots are smoothed by local regression using weighted linear least squares and a first-degree polynomial mode. The cumulative DAG and Rho signal is then presented as a histogram with a bin size of 10 degrees.

### Mathematical model of myosin regulation by PKC and ROCK

We assume that LPA is presented as a linear gradient and that the occupancy of LPA receptor follows a similar pattern. The receptor activates the PLC/PKC and Rho/ROCK pathways, the outputs of which are PKC and ROCK kinase activities. The key distinction between the two is that PKC activation is “polarized”, in the sense that there is intense activation at the front of the cell and little or none elsewhere, whereas ROCK activation is graded, similar to the receptor occupancy. We adopt the simple, steady-state models of diffusible intracellular messengers presented in Haugh and Schneider (Haugh and Schneider, 2006) to derive expressions for PKC and ROCK activities as a function of the spatial coordinate, *x* (*x* = 0 is the front of the cell, *x* = 1 is the rear). The following PKC function assumes a nonzero flux at *x* = 0 (mimicking the localized production at the front), satisfies no flux at *x* = 1, and is normalized by the value at *x* = 0.

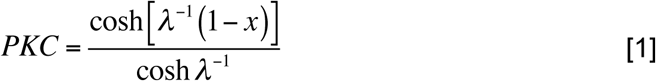

The pattern is determined by the balance of messenger diffusion and degradation, characterized by the dynamic length scale, *λ* (a value of *λ* = 0.1 was used; i.e., we assume that the average distance that a messenger molecule diffuses during its lifetime is 10% of the cell length). For *λ* << 1 as assumed, *PKC* ≈ *e*^−*x/λ*^. The following expression for ROCK assumes a linear source term with steepness *δ* (*δ* = 0.2 was used; i.e., assuming a 20% difference in production rate across the cell), satisfies no-flux conditions at both *x* = 0 and *x* = 1, and is normalized by the value at *x* = 0.

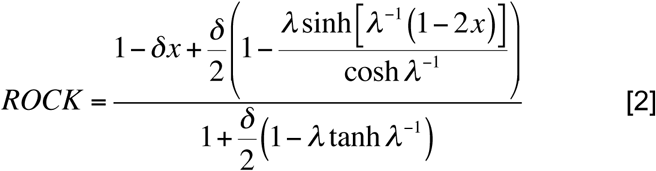

For simplicity, we assume the same value of the dynamic length scale, *λ* = 0.1; with *λ* << 1, the pattern of ROCK activity is approximately linear.

To conceptualize the effects of PKC and ROCK on myosin activity, we postulate that the two kinases phosphorylate MLC independently, but Ser1/2 phosphorylation by PKC ablates myosin activity regardless of Thr18/Ser19 phosphorylation status. We further assume that myosin diffusion is slow relative to the rates of MLC phosphorylation and dephosphorylation. Accordingly, we define the ratios of phosphorylated to unphosphorylated Thr18/Ser19 and Ser1/2 as *K*_18/19_ (1+ *αROCK*) and *K*_1/2_ *PKC*, respectively; values of *K*_18/19_ = 0.3, *α* = 10, and *K*_1/2_ = 10 were assumed (it suffices that *α* and *K*_1/2_ are >> 1). Based on these assumptions, the myosin activity as a fraction of the maximum is derived; the maximum activity corresponds to complete phosphorylation of Thr18/Ser19 and none on Ser1/2.

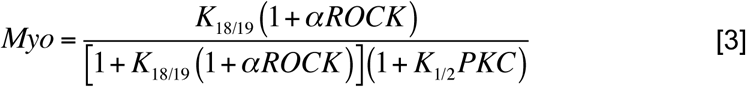

## SUPPLEMENTAL MATERIAL

**Supplemental Fig. 1:**
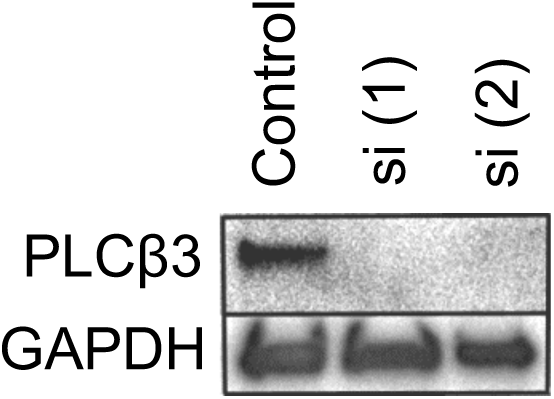
(relating to main Fig. 3D) Western blots show level of PLCβ3 depletion using two different siRNAs.

**TABLE S1:** The number of cells tracked for each experiment and corresponding velocity.

| | Vel ($\mu\text{m/hr}$) | n |
| --- | --- | --- |
| IA32 fibroblasts in LPA gradient | $17.86 \pm 1.09$ | 84 |
| IA32 fibroblasts + 10$\mu\text{M}$ Ki 16425 in LPA gradient | $16.42 \pm 1.72$ | 55 |
| IA32 fibroblasts in 10% serum gradient | $18.6 \pm 4.21$ | 45 |
| IA32 fibroblasts + 10$\mu\text{M}$ Ki 16425 in 10% serum gradient | $24.6 \pm 6.60$ | 46 |
| IA32 fibroblasts +150mM CK666 in LPA gradient | $24.3 \pm 5.1$ | 89 |
| Mouse dermal fibroblasts (pre-cre) in LPA gradient | $14.6 \pm 4.21$ | 48 |
| *Arpc2*<sup>-/-</sup> mouse dermal fibroblasts (post-cre) in LPA gradient | $9.51 \pm 1.93$ | 67 |
| IA32 fibroblasts + 15$\mu\text{M}$ Blebbistatin in LPA gradient | $24.1 \pm 8.2$ | 123 |
| Myosin IIA depleted IA32 fibroblasts in LPA gradients | $21.6 \pm 1.6$ | 52 |
| Myosin IIB depleted IA32 fibroblasts in LPA gradient | $23.6 \pm 2.31$ | 41 |
| *Arpc2*<sup>-/-</sup> mouse dermal fibroblasts+ 15 $\mu\text{M}$ blebbistatin in LPA gradient | $8.3 \pm 0.6$ | 44 |
| IA32 fibroblasts expressing myoRLC-GFP in LPA gradient | $12.04 \pm 1.1$ | 63 |
| IA32 fibroblasts expressing myoRLC(S1AS2A)-GFP in LPA gradient | $8.89 \pm 0.73$ | 78 |
| Mouse dermal fibroblasts (pre-cre) in LPA gradient -3D migration | $9.34 \pm 2.6$ | 34 |
| *Arpc2*<sup>-/-</sup> mouse dermal fibroblasts (post-cre) in LPA gradient-3D migration | $7.51 \pm 1.09$ | 42 |
| REF52 fibroblasts in LPA gradient-3D migration | $10.12 \pm 2.42$ | 64 |
| Myosin IIA depleted REF52 fibroblasts in LPA gradients – 3D migration | $13.08 \pm 3.52$ | 51 |
| Myosin IIB depleted REF52 fibroblasts in LPA gradient – 3D migration | 6.02 ± 0.85 | 47 |
| IA32 fibroblasts + 5mM PI3K inhibitor in LPA gradient | 24.1 ± 8.2 | 123 |
| PLCg1-null fibroblasts in LPA gradient | 10.03 ± 2.46 | 36 |
| PLCb3 depleted IA32 fibroblasts in LPA gradient | 11.30 ± 0.89 | 48 |
| IA32 fibroblasts expressing dominant negative PLCb3 in LPA gradient | 7.14 ± 1.62 | 31 |
| PKCα depleted IA32 fibroblasts in LPA gradient | 15.23 ± 3.64 | 38 |
| IA32 fibroblasts in PDGF gradient | 18.23 ± 2.14 | 54 |
| IA32 fibroblasts + 15 μM ROCK inhibitor in LPA gradient | 11.10 ± 1.18 | 51 |
| IA32 fibroblasts + 5 μg/ml RHO activator in PDGF gradient | 11.37 ± 1.10 | 81 |

## SUPPLEMENTAL MOVIE LEGENDS

**Movie 1:** IA32 fibroblast cells chemotaxing to a gradient of LPA. Images captured every 10 minutes with a 20X objective.

**Movie 2:** IA32 fibroblast cells exposed to a uniform concentration of LPA-R inhibitor, Ki 16425 (10μM) in gradient of LPA are unable to chemotax. Images captured every 10 minutes with a 20X objective.

**Movie 3:** Arpc2 null fibroblasts chemotaxing to a gradient of LPA. Images captured every 5 minutes with a 60X objective.

**Movie 4:** Arpc2 null fibroblasts migrate by extending filopodial protrusions in the direction of the LPA gradient. Images captured every 2 minutes with a 60X objective and 2X optical zoom.

**Movie 5:** MyoRLC-S1AS2A fibroblasts (fluorescent mutant cell in frame 1 top left corner) are unable to chemotax while control cells chemotax in an LPA gradient. Images captured every 10 minutes with a 20X objective.

**Movie 6:** Cells expressing RBD-GFP chemotaxing to a gradient of LPA. Images are acquired by Total Internal Reflection Fluorescence (TIRF) microscopy every 2 minutes for up to 6 hours. Profile of active Rho translocated to the membrane is observed using this technique.

**Movie 7:** Cells expressing PKCδ-GFP chemotaxing to a gradient of LPA. Images are acquired by Total Internal Reflection Fluorescence (TIRF) microscopy every 2 minutes for up to 6 hours. Profile of active DAG translocated to the membrane is observed using this technique.

**Movie 8:** Fibroblasts chemotax to an LPA gradient in the presence of Y compound. At time point 00:00:07, the drug is washed out and the cells regain typical fibroblast morphology and continue to chemotax with higher fidelity.

**Movie 9:** Fibroblasts chemotax to a PDGF gradient in the presence of Rho activator.

